# Comparative Genomics Exploration of Shared lncRNA-coding genes in Human and All Annotated Plant Genomes: Potential Implications for Cancer Research

**DOI:** 10.1101/2023.10.25.563922

**Authors:** Ligia Mateiu

## Abstract

Since their discovery in 1990, lncRNAs have been heavily investigated. It is already known that the genomes of plants and animals express large numbers of lncRNAs. In humans, many lncRNA are considered essential players in cancer biology, mainly acting as gene expression regulators in processes like cell proliferation, apoptosis, migration, invasion and stemness. In plants, lncRNAs are involved in vegetative growths, reproduction and stress responses. Yet, in the entirety of the animal and plant kingdoms, the functions and mechanisms of action for the majority of lncRNAs remain largely undiscovered, as do their orthology relationships. The poor conservation of the DNA sequences encoding for these transcripts is hindering the identification of lncRNAs orthologs especially among very distant evolutionary species. In this short study, using an uncomplicated, but creative bioinformatic approach, I searched for sequence homologies between human DNA and 175 annotated plant genomes from NCBI. Using stringent filtering, I found 20 human lncRNA-encoding genes overlapping plant genomic features, including lncRNAs. Evenmore, 3 of these human lncRNA-encoding genes with a potential plant ortholog are reported in multiple databases for lncRNA involved in cancers. This study opens the road for investigating the tumorigenesis in a deep homology context of lncRNAs.

## Introduction

Among the vast number of transcribed RNA molecules that reach the cytoplasm, only a minor fraction is further processed and becomes a functional protein in a certain tissue. The majority of transcripts are non-coding and they orchestrate many cellular functions, mainly gene expression. Over the past few decades, there has been a significant focus on a special category of non-coding RNAs, specifically long non-coding RNAs (lncRNAs), with more than 53,000 publications in PubMed to date. In humans, lncRNAs are considered essential players in cancer biology, mainly acting as gene expression regulators in processes like cell proliferation, apoptosis, migration, invasion and stemness ^1^. Thus, it is unsurprising that conducting an additional PubMed search for keywords “human AND lncRNA AND cancer” yields more than 22,000 results, including many databases and atlases for their functional annotation and characterization. However, there is still limited knowledge on their classification, their functions and also their conservation across evolutionary close and distant species ^2^. In a recent comparative genomics review ^3^, the authors summarize the challenges associated with identifying lncRNA orthologs. Among these challenges is the rapid evolution of lncRNAs, resulting in low sequence similarity even in closely related species. However, some syntenic lncRNAs were identified between human and lancelets by other means than sequence similarity analysis ^4^.

In this brief bioinformatic investigation, I conducted a comparative analysis of the DNA of all annotated plant genomes from NCBI (175) and the most recent human reference genome (ENSEMBL build 110). I achieved this using a cutting-edge computational tool specifically designed for processing Oxford Nanopore long reads, completing the analysis in an impressively short time frame, in a matter of a few hours. Through this method, I successfully identified instances where plant DNA exhibited significant sequence similarity with human lncRNA-encoding genes, observed across multiple plant species. Upon closer examination of these plant regions, it became evident that certain plant lncRNA-coding genes mapped sufficiently well to some human lncRNA-coding genes, according to NCBI genome annotation files. Intriguingly, a subset of human lncRNAs-coding genes on the short arm of chromosome 21 that exhibit potential plant orthologs were also found in databases dedicated to lncRNAs associated with cancer.

## Materials and Methods

**Fig. 1** provides a visual representation of the essential steps taken during the comparative analysis between the human reference genome (hg38 build version 110 from ENSEMBL) and the 175 annotated reference genomes of plant species retrieved from NCBI RefSeq. The DNA sequences and their latest annotations were programmatically obtained by following the links provided in the assembly_summary.txt file located on the NCBI FTP site (https://ftp.ncbi.nlm.nih.gov/genomes/refseq/plant/). The combined size of the compressed DNA sequence files (FASTA format) for 175 plant reference genomes, along with their corresponding gene structure information files (GTF format), was just under 52 gigabytes.

**Figure 1.**
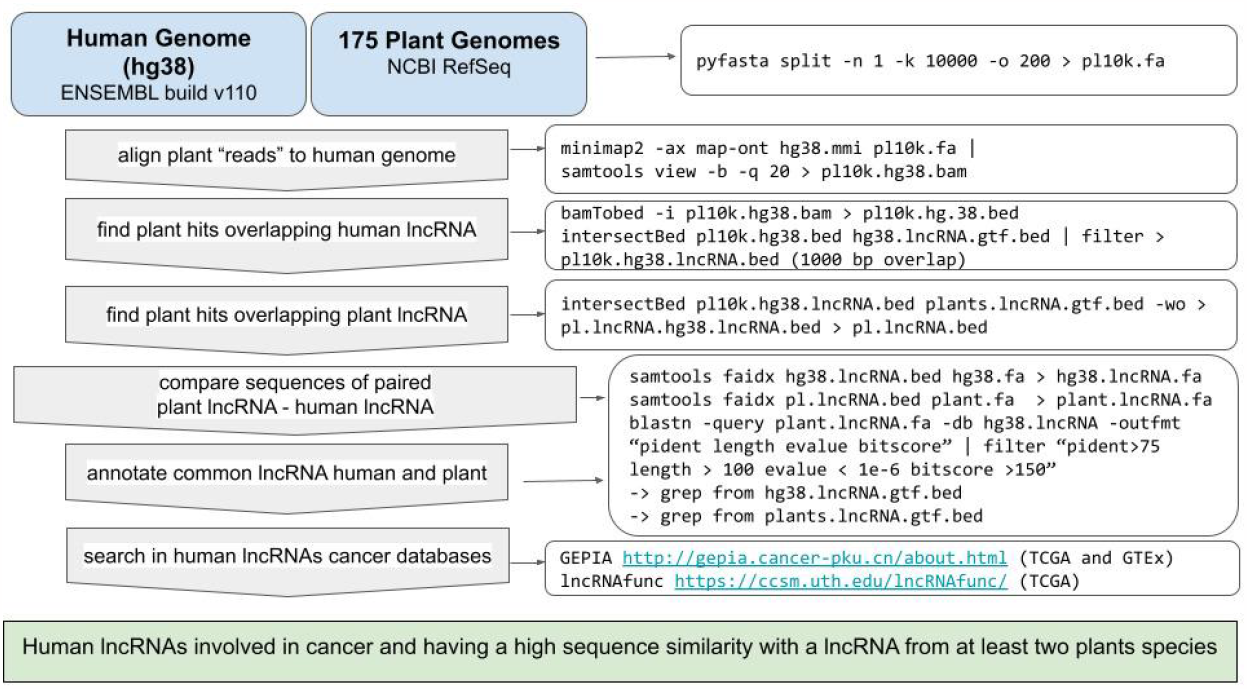
Illustration of the workflow employed in the comparative genomics analysis to identify shared lncRNAs in both human and plant genomes. The input data is presented within the blue boxes, while the bioinformatics tools, along with their respective parameters for data filtering, are shown on the right side.

The objective of this study is to identify sequence similarities between the human genome and all annotated plant genomes. Given the evolutionary divergence between humans and plants, direct genome-to-genome alignment is impractical. To tackle this, I put forward a straightforward solution: I divided each plant genome, arranged by chromosomes in FASTA format, into 10,000-base pair fragments (10k-mers) using the *pyfasta* bioinformatics tool ^5^, while preserving the genomic coordinates in the FASTA header. These fasta “reads” were then aligned to the human genome using *minimap2* (version 2.20-r1061) ^6^, a tool designed for long-read alignment, typically acquired from Oxford Nanopore sequencing. The method of indexing the reference genome and the approach to handling low-identity regions contributed to the attractiveness of this tool for the proposed comparative analysis. While the tool offers the flexibility to map different species, the substantial divergence between human and plant genomes posed a significant challenge in identifying extensive overlapping regions. To address this challenge, I adopted a workaround solution by treating the 10k-mer plant sequences as failed (bad quality) Oxford Nanopore long reads. For the subsequent analysis, I chose only the ‘reads’ mapped with a quality score greater than 20. The next goal was to identify regions of overlap with lncRNA genes in the human genome annotation file. In the current ENSEMBL genome build version (v110), out of a total of all 62,754 gene features, 18,866 were classified as having the lncRNA biotype.

Using *bedtools* ^7^, I intersected the 10k-mer coordinates of the plant genomes that mapped to human lncRNA genomic locations. Using only the plant segments with a minimum overlap of 1000 base pairs, I investigated whether they intersected with plant lncRNAs, as provided in the GTF (Gene Transfer Format) file for each respective organism. Next, I determined the exact sequence similarity for each pair of human and plant lncRNAs-coding genes by using the command-line *blastn* (*megablast)* tool ^8^ with stringent filtering (percent identity >75%, alignment length > 125bp, alignment E-value < 1e-6 and bitscore >150.

In the final stage of the analysis, I explored whether the human lncRNAs with potential plant orthologs are functional, specifically examining whether they had been reported as differentially expressed in tumor versus normal tissues. I selected two recently published and popular knowledge bases of lncRNA function in human cancer lncRNAfunc ^9^ and GEPIA2 ^10^. LncRNAfunc serves as a comprehensive resource for annotating the functions of almost 16,000 lncRNAs across 33 different types of cancers in TCGA (The Cancer Genome Atlas), containing a wide array of information such as interactions, gene expression modifications, alternative splicing, and more. GEPIA2, also a web-based platform, offers extensive functional information for both coding and non-coding transcripts across 84 cancer subtypes, utilizing TCGA and GTEx (Genotype-Tissue Expression) data.

## Results

The comparative study involving 175 reference plant genomes and human DNA started with the segmentation of plant genomes into 10,000 base-pair segments (10k-mers), with a 200 base-pair overlap between adjacent tiles using the bioinformatics tool *pyfasta* (the results obtained at every stage of the analysis are summarized in **Fig. 2)**.

**Figure 2.**
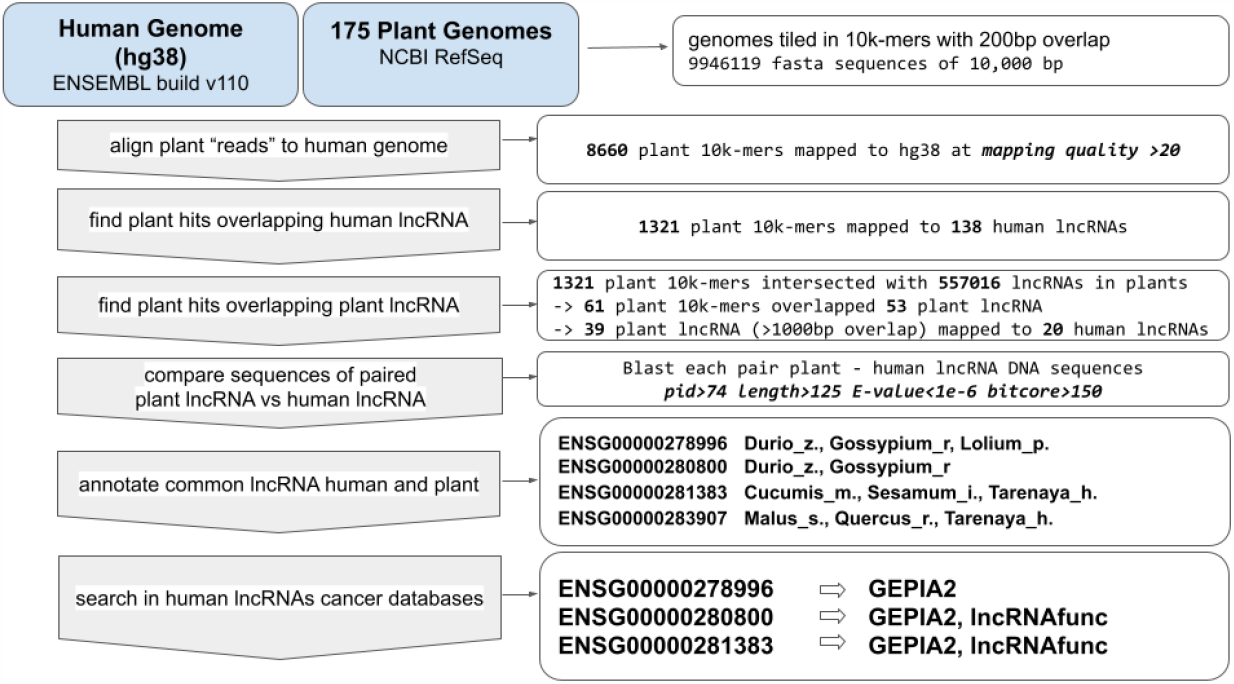
Results of the comparative genomics analysis, step by step. The figure illustrates the count of plant and human sequences obtained at each stage of the analysis. The ENSG numbers represent the human long non-coding RNAs (lncRNAs), identifiers in ENSEMBL, for which there is a “corresponding” lncRNA in a minimum of two plant species.

Subsequently, the approximately 10 million plant 10k-mers were mapped to the human reference genome (hg38) using *minimap2*. Thus, I retrieved 8,660 plant 10k-mers that exhibited a minimum mapping quality of 20 when aligned to the human genome. The 10k-mers mapping to multiple locations within the human genome were excluded. Employing *bedtools*, I identified 1,321 plant 10k-mers that overlapped with 138 distinct human lncRNAs and I proceeded with the analysis by selecting the 10k-mers that had overlaps exceeding 1000 base pairs with a human lncRNA-coding gene. The count of interesting locations decreased drastically when I narrowed it down to the 10k-mers that overlapped with a plant lncRNA, as outlined in the plant genome annotation files. More precisely, the count of 10k-mers trimmed down to 61, and this number further reduced to just 39 after implementing the 1000bp overlap filtering criteria. The DNA sequences of these plant lncRNAs mapped to 20 human lncRNAs to a certain extent. The subsequent phase of the analysis was dedicated to this reduced subset, for which I extracted the DNA sequences corresponding to their genomic coordinates and measured the homology between them. I filtered the *blastn* output to retain pairs with a percent identity greater than 75%, a query vs. sequence alignment size exceeding 125, an E-value lower than 1e-6, and a bitscore over 125.

Ultimately, only four human lncRNA-coding genes remained, each of which had a corresponding plant lncRNA-coding gene in at least two distinct plant species. This limited count is not unexpected, given the significant divergence in DNA sequences between humans and plants, coupled with the stringent filtering criteria that excluded micro-homologies with shorter lncRNAs. An overview of these human lncRNAs is presented in **Fig. 3**.

**Figure 3.**
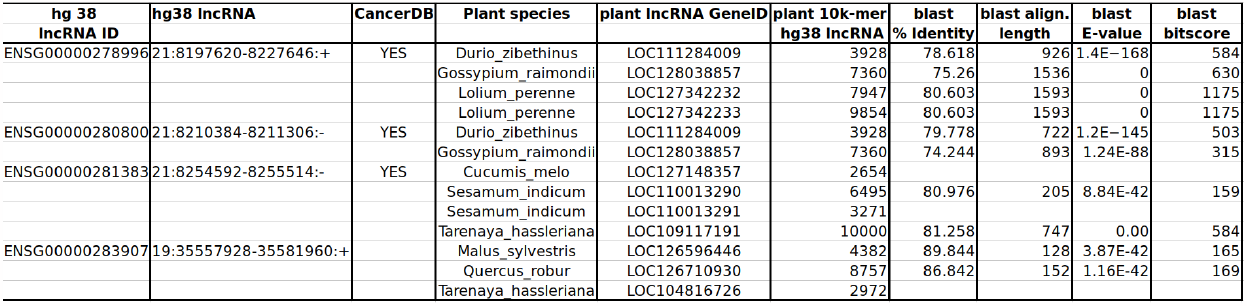
Overview of the human lncRNA-coding genes exhibiting substantial sequence similarity with lncRNA-coding genes found in a minimum of two plant species. The comprehensive overview shows the ENSEMBL identifiers; genomic coordinates in the hg38 v110 build; the presence of human lncRNAs in cancer databases GEPIA2 and longRNAfunc; plant species names and genomic coordinates for the selected plant lncRNAs; the extent of overlap between the initial plant 10k-mers and human lncRNAs; blast percent identity for the human - plant lncRNA pair, blast alignment length, and blast E-value and bitscore for evaluating the quality of the sequence comparison.

ENSG00000278996, ENSG00000280800, ENSG00000281383 are all novel transcripts located on chromosome 21. Perhaps this finding is not entirely unexpected, as chromosome 21 is known for its high density of lncRNA-encoding genes per megabase ^11^. Nevertheless, it’s quite fascinating that there is a significant sequence similarity spanning approximately 1200 nucleotides with lncRNAs in specific annotated plant species: Durio_zibethinus, Gossypium_raimondii, Lolium_perenne, Cucumis_melo, Sesamum_indicum, Tarenaya_hassleriana. For a more detailed examination of how the plant sequences aligned with two human lncRNA-encoding genes, see **Fig. 4**.

**Figure 4.**
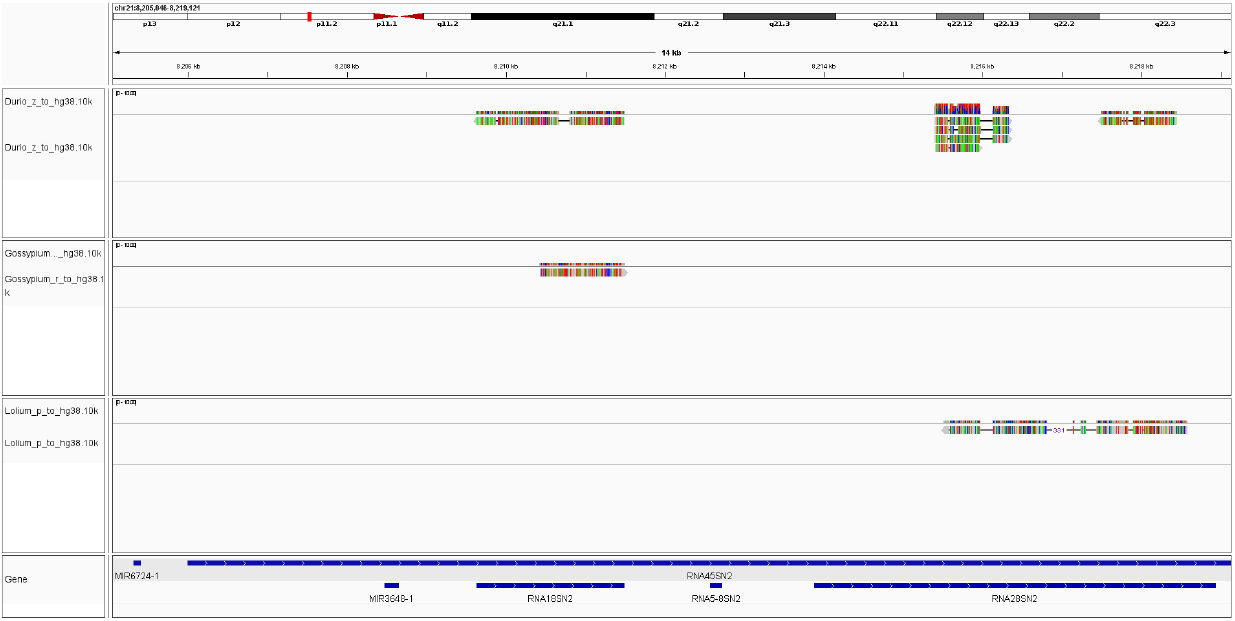
IGV snapshot of the human genomic region chr21:8,205,046 - 8,219,121. In this region, 10k-mers from Durio zibethinus, Gossypium raidomdii and Lolium perenne were mapped by minimap2 at mapping quality over 20. ENSG00000278996 and ENSG00000280800 are located within these coordinates.

ENSG00000283907 is also a lncRNA characterized as a novel transcript, antisense to ATP4A. The gene responsible for encoding this transcript has approximately 125 nucleotides overlapping with lncRNA genes in Malus_sylvestris, Quercus_robur, Tarenaya_hassleriana.

Given that lncRNAs are known to play a role in regulating a wide range of cellular processes, I opted to investigate whether the expression of these four lncRNAs is perturbed in cancers. I used two popular web-based platforms: GEPIA2 and lncRNAfunc. All three lncRNAs located on chromosome 21 were documented in GEPIA2 as having modified expression in several cancer types, with the predominant trend being up-regulation. ENSG00000280800 and ENSG00000281383 were also reported in lncRNAfunc. The links to these databases are available in the **Suppl. Table 1**.

**Suppl. Table 1. Mapped 10k-mer plant DNA sequences overlapping 20 human lncRNA-encoding genes**. The table contains all the plant 10k-mers that mapped to some extent to a human lncRNA-encoding gene.

Although the plant lncRNAs found in this concise study are uncharacterized, this could stand out as a relevant discovery: the prospect of exploring the plant genomes for addressing human cancer-related issues. I’m alluding to this because research into the potential roles and regulatory mechanisms of lncRNAs in plant development is gaining significant momentum ^12^.

The bash scripts and all the results are available in the github repository https://github.com/ligiamateiu/pair_plant_human_lncRNA/

## Discussion

In this concise bioinformatic study, I found 4 human lncRNA-encoding genes that show sufficient sequence similarity with lncRNA-encoding genes from several plant species. This study serves as a proof of principle, suggesting the possibility of shared ancestral and potentially functional lncRNAs between humans and plants. Their location on the short arm of chromosome 21 adds an even more intriguing dimension, as this region is frequently scrutinized in studies focused on evolutionary conservation ^13–15^. Considering that the lncRNAs encoded by these potentially shared human and plant genes are expressed and demonstrated to be functional in cancer, it raises the intriguing questions: Could the origins of cancers be linked with the lncRNAs evolution? How old are the cancer genes?

## Abbreviations

lncRNA: long non-coding RNA
GTF: Gene Transfer Format
TCGA: The Cancer Genome Atlas
GTEx: Genotype-Tissue Expression

## Notes

https://github.com/ligiamateiu/pair_plant_human_lncRNA

### Competing Interest Statement

The authors have declared no competing interest.

